# The Influence of Public Health Faculty on College and University Plans during the COVID-19 Pandemic

**DOI:** 10.1101/2021.06.23.449583

**Authors:** David A. Johnson, Meredith Cahill, Sara Choate, David Roelfs, Sarah E. Walsh

## Abstract

The purpose of this study was to determine whether the institutional presence of public health faculty within colleges and universities influenced operational plans for the fall semester of 2020. Using cross-sectional data collected by the College Crisis Initiative of Davidson College, six levels of instructional modalities (ranked from least to most restrictive) were compared between Council on Education of Public Health (CEPH)-accredited and non-CEPH-accredited 4-year institutions. Institutions with CEPH-accredited schools and programs were more likely to select some restrictive teaching modalities: 63.8% more likely to use hybrid/hyflex or more restrictive and 66.9% more likely to be primarily online (with some in person) or more restrictive. However, having CEPH-accredited programs did not push institutions to the most restrictive modalities. COVID-19 cases in county, enrollment, and political affiliation of the state governor were also found to influence instructional modality selection. While any ecological study has certain limitations, this study demonstrates that college and university fall plans appear to have been influenced by the presence of CEPH-accredited schools and programs of public health, and/or the input of their faculty. The influence of relevant faculty expertise on institutional decision-making can help inform college and university responses to future crises.

## Introduction

In spring of 2020, higher education in the U.S. was faced with a problem not experienced in 100 years – deciding how to respond and operate during a global pandemic. What began in the spring semester with abrupt disruptions, led to the realization that the COVID-19 crisis would continue into the fall. Over the summer, virtually every college and university engaged in dialogue and strategic planning to formulate their response. In the U.S., this was reflected in a wide range of governance patterns informed by varying influences and controls. While the shared governance methodology provides structure through which leadership may meet its educational responsibilities, the COVID-19 crisis presented a new and formidable challenge – inserting public health as a primary focus in higher education’s collective decision-making model (Laforge, 2020).

As institutions of higher education considered the best response to address the needs of stakeholders, the global pandemic created three simultaneous existential crises— the threat to the population’s health, economic stability, and the unknown future of higher education (Widmer et al., 2020). To plan responsibly, Widmer et al^2^ argue that each institution, with its unique set of circumstances and resources, would need to consider its own equation of metrics, including; location, rate of infection based on its geography, testing capacity, availability of personal protective equipment, capacity to quarantine infected students living on campus, and other factors. Fundamentally, postsecondary institutions are complex organizations that act in the often-competing interests of their educational mission and the bottom line (Widmer et al., 2020).

Prior to the COVID-19 pandemic, public colleges and universities were grappling with declining state funding, which never fully recovered from the austerity measures of the 2008 Great Recession (Mitchell et al., 2018). This has led to an increased reliance on tuition dollars and other revenue-generating activities, such as residence halls and university events (Nadworny, 2020), at a time when there were fewer students available to pay tuition (*Current Term Enrollment Estimates*, 2020). Indeed, colleges and universities which were more dependent on revenue from auxiliary services provided to on-campus students were more likely to open for inperson instruction (Klinenberg & Startz, 2021). Amid these economic threats related to decreased enrollment, as well as the global pandemic’s damaging effect on national, state, and local economies, there was backlash from students and families who were angry for being charged the same or more for online learning in lieu of in-person courses in the spring (Anderson, 2020).

Aside from economics, other principles likely guided fall decision making in higher education. Organizational theory stresses the strong pressures organizations face to conform to emergent norms - i.e., become isomorphic (DiMaggio & Powell, 1983) - and to otherwise be bound to the dominant “institutional logic” in an industry (Thornton et al., 2012). Additionally, organizational decision making is conditioned by the feedback received from stakeholders, as well as the resources/knowledge available to seek solutions to problems (Greve, 2003). Previous explorations of fall reopening plans have concluded that colleges and universities were influenced by state and local politics as much, if not more than, institutional characteristics (Collier et al., 2021; Felson & Adamczyk, 2021). This offers insight into decision-making processes of institutions, given their collective limitations of time to design a reopening plan and knowledge about how to operate under a pandemic. Given these competing forces, it was unclear whether public health would have a meaningful voice in campus responses to COVID-19.

Public health influence would likely be shaped by the Precautionary Principle, a foundational tenet within both public health training and practice. The Precautionary Principle includes the following components: (1) taking preventive action in the face of uncertainty, (2) shifting the burden of proof to proponents of an activity, (3) exploring a wide range of alternatives to potentially harmful actions, and (4) increasing public participation in decision making (Kriebel & Tickner, 2001). Much of the uncertainty of the fall semester has given way to the consequences of instructional modality decisions. Studies have linked the return of students to campus to COVID-19 surges both on campus (Harris, 2020) and in the surrounding community (Leidner, 2021). While these studies highlight the stakes of these fall plans, it is necessary to understand how colleges and universities made these decisions in the face of uncertainty, with only the information available at the time.

The COVID-19 pandemic of 2020 has created a new opportunity for the field of public health and its professionals to help guide coordinated efforts at national, state, local, and organizational levels. Within institutions of higher education, principles of shared governance necessitate consultation with public health faculty when such professionals are present within the academy. Like all faculty, their research and instructional efforts are critical to a university’s business model, but moreover, public health faculty have specific subject matter expertise relevant to the crisis at hand and are uniquely qualified to help navigate a global pandemic. Public health faculty influence was selected for this study because of the disciplinary focus on prevention, population health, and community spread of disease, rather than the treatment of individuals and outcome-focused expertise from medicine, nursing and other clinical professions.

A better understanding of the decision-making processes employed throughout higher education can inform the continued response to the COVID-19 pandemic, as well as preparedness efforts for future crises. This research seeks to better understand the factors related to the development of fall plans and instructional strategies and factors influencing the final decisions. Included in these potential factors is an examination of whether the presence of academic public health professionals, within these colleges and universities at accredited schools and programs of public health, had bearing on the selected modalities for fall plans.

## Materials and Methods

Cross-sectional data were derived for 3,036 institutions of post-secondary education through the compilation of data from multiple sources. Data on university course instruction modality (for fall 2020) and fall enrollment (for fall 2018) were obtained from the College Crisis Initiative (C2i). C2i was created by Davidson College to track university responses to COVID-19, which also provided a 7-day rolling average for county-specific incidence rates, dated to August 21^st^, 2020. Instruction modality at the beginning of the fall 2020 semester was coded by C2i into fourteen separate modalities, but for the purpose of clarity, we used the Chronicle of Higher Education’s (*Here’s Our List of Colleges’ Reopening Models*, 2020) modality designation that collapses many of the instructional strategies into a 6-point ordinal scale from least restrictive to most restrictive: 1) fully in-person instruction; 2) primarily in-person; 3) hybrid/hyflex, professor’s choice, simultaneous teaching, some variety of methods and, or non-specific plan; 4) primarily online; 5) fully online with some students on campus; 6) fully online instruction with no students on campus. We removed any institutions listed as Closed, No COVID Mentions, TBD/Undetermined, and Other from the analysis.

Colleges and universities were denoted as having an institutional presence of public health faculty based on established schools of public health, public health programs, or standalone public health baccalaureate programs accredited by the Council on Education for Public Health (CEPH). Though public health is taught in unaccredited settings as well, the analysis focused on CEPH institutions due to the curricular standards and quality conferred through the accreditation process. Additionally, we posited that university administrators may be more aware of faculty public health expertise if they have allocated the staff, time, and other resources necessary for the accreditation process. Current state governor’s party affiliation was derived from Ballotpedia (*List of Governors of the American States*, n.d.) and the presence of an American Association of University Professors (AAUP) chapter at each institution was derived from the AAUP official list located on its website (*Finda Chapter / AAUP*, n.d.).

Of the 3,036 colleges and universities in the C2i dataset, we focused our analyses on the 2,001 from which students may receive a 4-year degree. Of these, 159 were excluded from the analysis due to missing data on instructional modality and an additional 78 were excluded due to missing data on enrollment. The analyses were therefore based on the 1,764 four-year colleges and universities for which we had full data, or 86.6% of the total number of listed 4-year institutions.

The dependent variable (DV) used in the analysis was the six-level modality restriction scale described above. Five independent variables (IVs) were used in the analysis. First, we included a binary variable for whether a college or university has a CEPH-accredited program on campus. For the purposes of the analysis, satellite campuses are not counted as having a CEPH-accredited program since it could solely be housed on the main college or university campus. Second, we included a count variable for fall 2018 student enrollment. Third, we included a rate variable for the number of COVID-19 cases per 100,000 population in the county in which the college or university is located. Fourth, we included a binary variable for whether a college or university has an AAUP chapter. Finally, we included a binary variable for the political party affiliation (Democrat or Republican) of the governor of the state in which a college or university is located. Mayor was substituted for governor in Washington D.C.

The data were analyzed using a mixed-effects ordered logistic regression model (using the meologit command in STATA 15.1) in order to account for clustering in the data by state (which have jurisdiction over educational policies to which all colleges or universities within the state must adhere). The data is therefore analyzed using a two-level regression equation, full detail found in Appendix (Equation A1).

The regression model was checked for multicollinearity and violations of the linearity and parallel lines assumptions. Multicollinearity was checked using Variance Inflation Factors. The model was checked for violations of the linearity assumption for the enrollment and COVID-19 rate variables using likelihood ratio tests comparing baseline models (linear predictors) against non-linear models using a restricted cubic splines transformation with 3 knots placed at the default percentiles (Harrell, 2001). Violations of the parallel lines (proportional odds) assumption were checked by comparing the results from the ordered logistic model to those of the five underlying binary logistic models. Based on the results of these tests, we modeled the COVID-19 rate using restricted cubic splines and included results of the underlying binary models as well as the ordered logistic model. It is important to note that, because this dataset is not from a sample, the statistics generated are population observations and not estimates (inference testing tools such as p-values and confidence intervals are therefore not appropriate, though we report confidence intervals in Table A1 in the Appendix along with an explanation for what confidence intervals mean in the context of population data).

## Results

Table 1 shows the prevalence of the six instruction modalities among the 1,764 4-year institutions included in the analysis, as well as the prevalence of CEPH-accredited programs, Republican governance, and AAUP chapters. The proportion of instruction modalities was 4.7% fully in-person; 29.1% primarily in-person; 25.9% hybrid/hyflex, professor’s choice, simultaneous teaching, some variety of methods, non-specific plan; 27.6% primarily online; 4.3% fully online, some students on campus; and 8.5% fully online, no students on campus. CEPH-accredited programs were present at 9.9% of the institutions within the analysis, or 182 in total. Fall enrollment varied greatly between colleges/universities, ranging from 18 to 121,437 students (mean = 6,748). The COVID-19 rate in the county also varied, ranging from 47 to 8993 cases per 100,000 population (mean = 1645.2). Only 15.7% of schools had an AAUP chapter. 53.2% of schools were located in a state where the governor’s political affiliation was Democratic (46.8% Republican).

**Table 1.** Descriptive Statistics^1^

| *Variables* | *Range* | *Median/Mean/%* | *Standard Deviation* |
| --- | --- | --- | --- |
| Modality restriction scale (levels shown below)<sup>2</sup> | 1 – 6 | 3 (median) |  |
| 1 = Fully in person |  | 4.7% |  |
| 2 = Primarily in person, some online courses |  | 29.1% |  |
| 3 = Hybrid/Hyflex teaching/Professor's choice/Simultaneous teaching/Some variety of methods, non-specific plan |  | 25.9% |  |
| 4 = Primarily online, some in person |  | 27.6% |  |
| 5 = Fully online, some students on campus |  | 4.3% |  |
| 6 = Fully online, no students on campus |  | 8.5% |  |
| Presence of CEPH school or program | 0 – 1 |  |  |
| No (reference group) |  | 90.1% |  |
| Yes |  | 9.9% |  |
| Fall 2018 enrollment<sup>3</sup> | 18 – 121,437 | 6,747.6 (mean) | 10,318.5 |
| COVID-19 cases per 100,000 in county<sup>4</sup> | 47 – 8993 | 1,645.2 (mean) | 980.2 |
| Presence of AAUP chapter | 0 – 1 |  |  |
| No (reference group) |  | 84.3% |  |
| Yes<sup>5</sup> |  | 15.7% |  |
| Political party of the state's governor | 0 – 1 |  |  |
| Democrat (reference group) |  | 53.2% |  |
| Republican |  | 46.8% |  |
| <sup>1</sup> n = 1764 four-year colleges/universities; 237 colleges/universities excluded from the analysis due to missing data on instructional modality (159 cases) and/or enrollment (78 additional cases) |  |  |  |
| <sup>2</sup> Dependent variable (DV) for the analysis |  |  |  |
| <sup>3</sup> Variable divided by 10,000 for the analysis |  |  |  |
| <sup>4</sup> Preliminary analyses showed the association with the DV to be non-linear; variable analyzed in the regression using the restricted cubic splines method |  |  |  |
| <sup>5</sup> n=182 four-year colleges/universities with CEPH-accredited programs out of 217; 35 accredited colleges/universities excluded from the analysis due to missing data on instructional modality |  |  |  |

The interpretation of the regression results hinges on the recognition that we are often examining associations with the DV for individual thresholds on that scale. In total, there are five thresholds: (1) DV levels 2-6 vs. 1 (having at least some classes online vs. being fully in person); (2) DV levels 3-6 vs. 1-2 (using hybrid/hyflex courses or something more restrictive vs. anything less restrictive); (3) DV levels 4-6 vs. 1-3 (using primarily-online instruction or something more restrictive vs. anything less restrictive); (4) DV levels 5-6 vs. 1-4 (fully online courses, with or without students on campus vs. anything less restrictive); and (5) DV levels 6 vs. 1-5 (fully online courses with no students on campus vs. anything else). For brevity, when referencing results for particular thresholds, we refer to these, respectively, as (1) Any restriction; (2) Hybrid+; (3) Primarily-online+; (4) Fully-online+; and (5) Fully-online-closed.

The logistic regression results (odds ratios) are reported in Table 2. Overall, the presence of a CEPH-accredited school or program was associated with a 34.4% increase in the odds of being at a higher lever on the modality restriction scale. However, as shown in Figure 1, the likelihood of choosing an instruction modality and the magnitude of influence varied by level of restriction.

**Figure 1:**
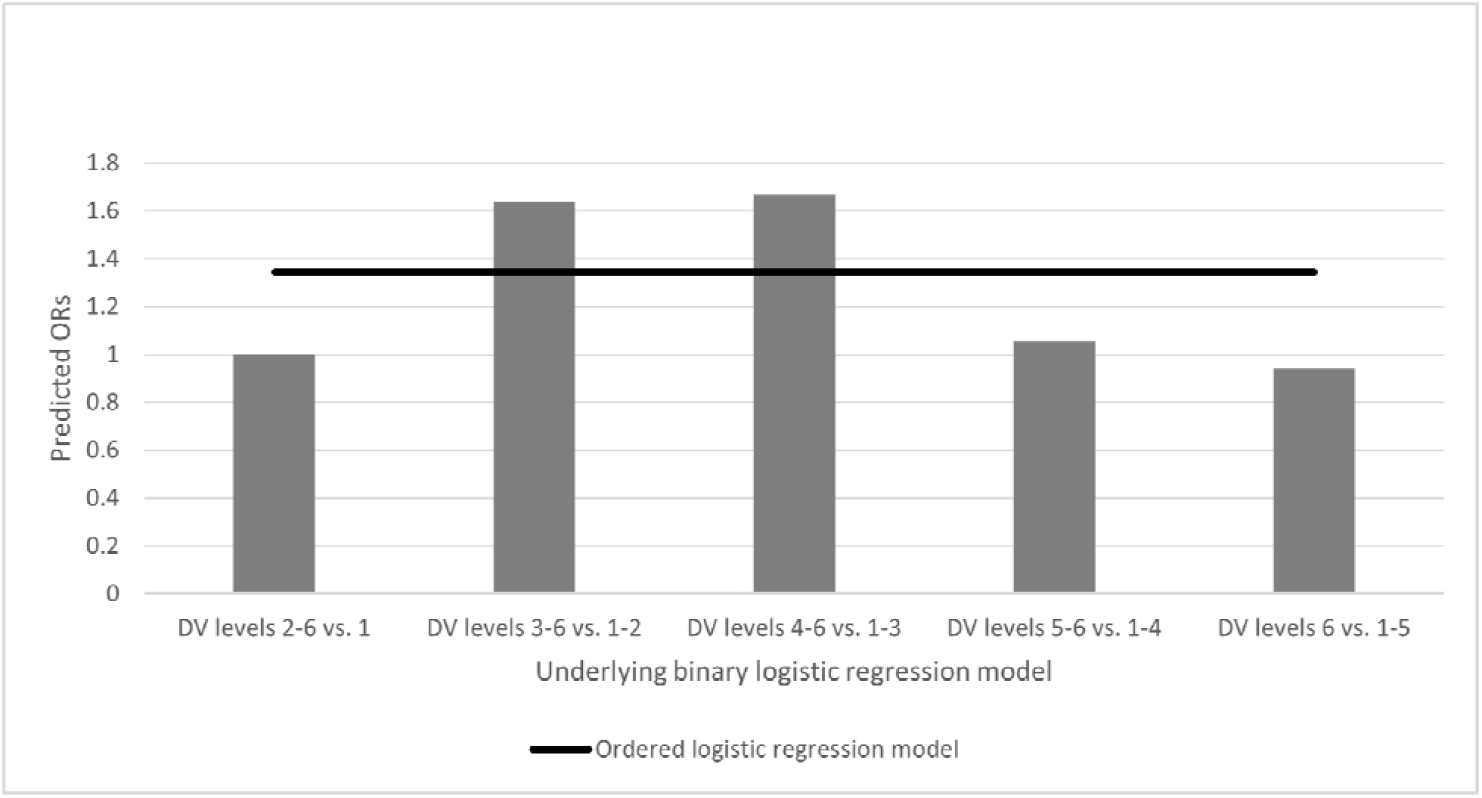
Predicted odds ratios for having a CEPH program, by regression model

**Table 2.** Mixed effect logistic regression models predicting the level of restriction on teaching modalities^1,2^

|  | *Full ordered logistic model* <sup>3</sup> | *Underlying binary logistic models* |  |  |  |  |
| --- | --- | --- | --- | --- | --- | --- |
|  |  | *DV levels 2-6 vs. 1* | *DV levels 3-6 vs. 1-2* | *DV levels 4-6 vs. 1-3* | *DV levels 5-6 vs. 1-4* | *DV levels 6 vs. 1-5* |
| **Institution-level portion of the model** |  |  |  |  |  |  |
| Presence of CEPH school or program | 1.3444 | 1.0040 | 1.6380 | 1.6690 | 1.0546 | 0.9422 |
| Fall 2018 enrollment (in 10000s) | 1.1263 | 6.4913 | 1.2445 | 1.1678 | 0.8703 | 0.6565 |
| COVID-19 cases per 100,000 in county |  |  |  |  |  |  |
| Linear effect | 1.0007 | 1.0004 | 1.0005 | 1.0005 | 1.0010 | 1.0011 |
| Non-linear effect <sup>4</sup> | 0.9994 | 0.9995 | 0.9995 | 0.9996 | 0.9990 | 0.9990 |
| Presence of AAUP chapter | 1.1257 | 2.9744 | 1.6484 | 1.0772 | 0.5230 | 0.4042 |
| Republican governor | 0.4482 | 0.3362 | 0.4802 | 0.4143 | 0.6469 | 0.6939 |
| Intercepts |  |  |  |  |  |  |
| DV levels 2-6 vs. 1 | 15.5720 | 15.6264 |  |  |  |  |
| DV levels 3-6 vs. 1-2 | 1.2375 |  | 1.1760 |  |  |  |
| DV levels 4-6 vs. 1-3 | 0.3645 |  |  | 0.4481 |  |  |
| DV levels 5-6 vs. 1-4 | 0.0649 |  |  |  | 0.0537 |  |
| DV levels 6 vs. 1-5 | 0.0400 |  |  |  |  | 0.0340 |
| **State-level portion of the model** |  |  |  |  |  |  |
| Intercept (variance) <sup>5</sup> | 0.5113 | 0.7328 | 0.5159 | 0.4741 | 0.4288 | 0.3948 |
| Republican governor (variance) <sup>6</sup> | 4.27e-31 | 1.07e-32 | 2.56e-37 | 3.12e-35 | 6.28e-34 | 1.01e-33 |
| **Model Log likelihood** | -2617.8121 | -288.9465 | -1034.1719 | -1091.1431 | -635.5080 | -482.6002 |

<sup>1</sup> Dependent variable (DV) is a 1-6 modality restriction scale, where 1 = “Fully in person”; 2 = “Primarily in person, some online courses”; 3 = “Hybrid/Hyflex teaching/Professor’s choice/Simultaneous teaching/Some variety of methods, non-specific plan”; 4 = “Primarily online, some in person”; 5 = “Fully online, some students on campus”; 6 = “Fully online, no students on campus”
<sup>2</sup> n=1764 four-year colleges/universities for all analyses; all numbers shown are odds ratios (where odds refers to the odds of being above a given threshold on the 1-6 modality restriction scale); all models computed using a mixed effects model with random intercepts by state and random slopes for the republican governor variable (the only other state-level variable in the model)
<sup>3</sup> The full ordered logistic model failed to meet the parallel lines assumption; underlying binary logistic models show where coefficients for the IVs vary at different thresholds/cut points on the DV
<sup>4</sup> Non-linear variable calculated using the restricted cubic splines method; see Figure 2 for the interpretation of the COVID-19 rate’s effect
<sup>5</sup> Number shown is the variance of the log odds; the results shown indicate a significant amount of variation in the intercept between states
<sup>6</sup> Number shown is the variance of the log odds; the results shown indicate no significant variation in the effect of the governor’s political party between states

The effects for institutional presence of a CEPH-accredited school or program appear concentrated at two thresholds: the decision to opt for Hybrid+ (63.8% higher odds versus less restrictive modalities) and the decision to opt for Primarily-online+ (66.9% higher odds versus less restrictive modalities). CEPH-accreditation was not as influential on the decision to employ Fully-online+. The presence of a CEPH-accredited program increased the odds of adopting a Fully-online+ by 5.4% and decreased the odds (by 5.8%) of selecting the most restrictive instructional method, Fully-online-closed. The presence of a CEPH-accredited school or program did not show an effect on universities’ decision to forgo fully in-person instruction.

Figure 2 shows the relationship between COVID-19 county incidence rate and the odds of choosing various levels of instructional modalities. At the median COVID-19 rate (approximately 1,500 cases), the odds of being at a higher lever on the modality restriction scale were 141.6% higher than they would be in the absence of COVID-19 cases. However, the odds varied across the DV scale, with a median infection rate leading to a 64.0% higher odds of having Any restriction; a 106.9% higher odds of being Hybrid+, a 102.7% higher odds of being Primarily-online+, a 295.7% higher odds of being Fully-online+, and a 338.2% higher odds of being Fully-online-closed. The influence of COVID-19 incidence rate on modality choice was greatest between 2,000 to 2,500 cases per 100,000 county population regardless of the threshold examined for the DV. After the rate of new cases surpassed 2,500 cases per 100,000 county population, the influence of incident rate on modality choice steadily declined for all restriction levels besides the decision to adopt a Primarily-online+ plan (though it is important to note that 92.4% of all schools were in counties with fewer than 3,000 cases per 100,000 population).

**Figure 2:**
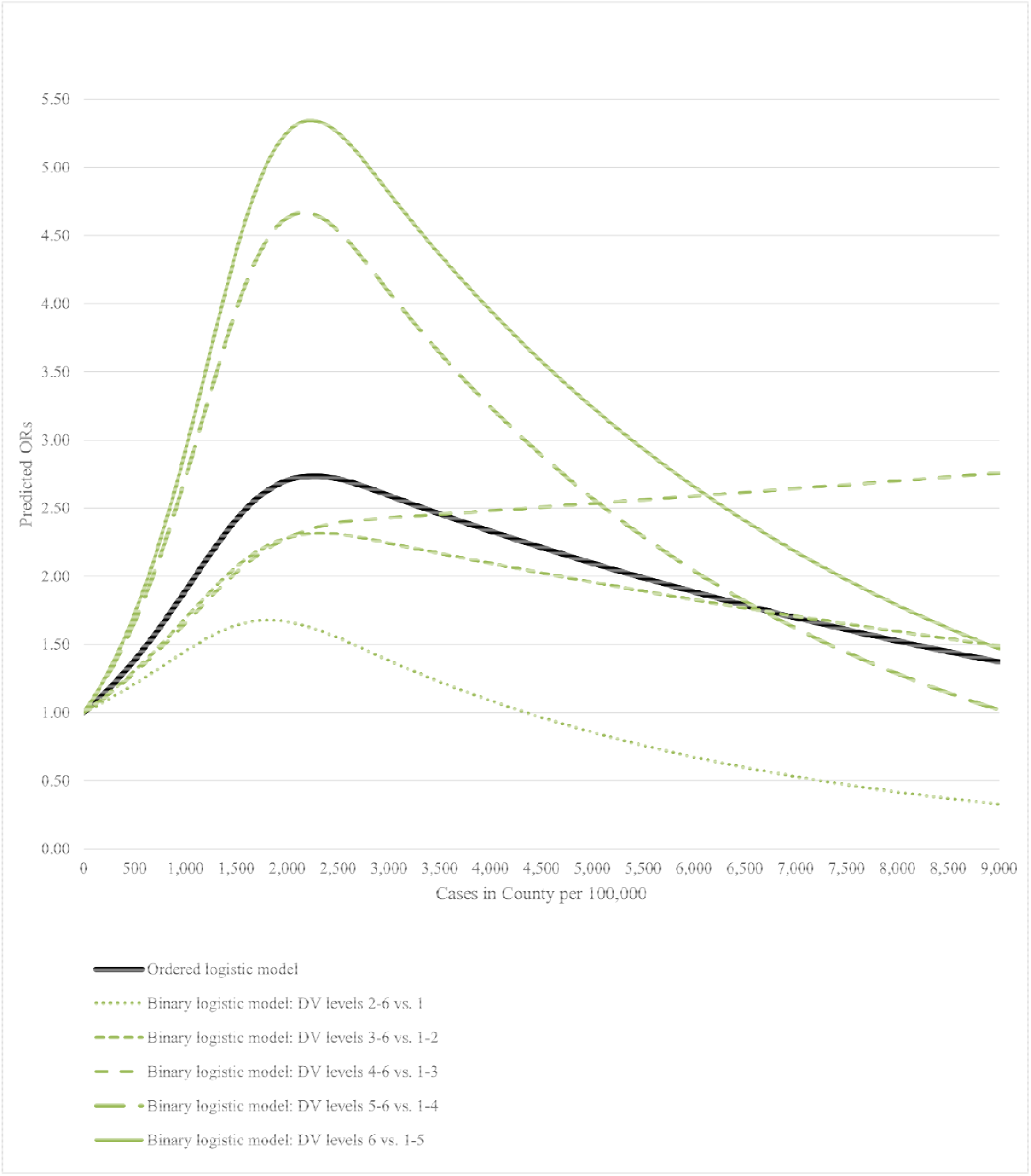
Predicted odds ratio by the COVID-19 incidence rate in the county

The association of fall enrollment with instructional modality followed a similar pattern to the COVID-19 incident rate. Overall, each additional 10,000 students at a school was associated with a 12.6% increase in the odds of being at a higher lever on the modality restriction scale, but the odds varied across the scale. We observed a very large increase in the odds for having Any restriction (549.1%), as well as increased odds of Hybrid+ (24.5%) and Primarily-online+ (16.8%). In terms of higher levels of restriction, each additional 10,000 students was associated with a 13.0% lower odds of being Fully-online+ and a 34.3% lower odds of being Fully-online-closed.

The presence of an AAUP chapter was associated with an overall 12.6% increase in the odds of being at a higher lever on the modality restriction scale. The association varied widely across the scale, with very strong increases in the odds of having relatively mild restrictions (197.4% higher odds of Any restriction; 64.8% Hybrid+; 7.7% Primarily-online+). In terms of higher levels of restriction, having an AAUP chapter was associated with a 47.7% lower odds of being Fully-online+ and a 59.6% lower odds of being Fully-online-closed.

Overall, we observed that schools in states with a Republican governor were 55.2% less likely to be at a higher lever on the modality restriction scale. In fact, at all DV thresholds, Republican governance decreased the likelihood of adopting more restrictive instructional modalities (Table 2). Schools in Republican governed states were 66.4% more likely to choose to teach courses fully in person and 58.6% less likely to opt for modalities more restrictive than Hybrid+.

## Discussion

The institutional presence of a CEPH-accredited school or program was found to be associated with college and university modality selection for fall 2020 when controlling for both enrollment and incidence of COVID-19 at the county level, as well as political affiliation of the governor at the state level. Institutions with these public health professionals as members of the academy were less likely to embrace a fully or primarily in-person approach, arguably the riskiest modalities given the threat and reality of the COVID-19 pandemic. However, having this institutional presence of public health professionals did not push institutions to the most conservative modalities of fully-online with students (Fully-online+) or fully-online without students on campus (Fully-online-closed). Instead, it appears that institutions with CEPH-accredited schools and programs opted for the middle approaches of either a hybrid/hyflex (Hybrid+) or primarily online modality (Primarily-online+).

The study did not intend to explore the COVID-19 infection data in a nonlinear method, but with this methodologically-necessary approach came the finding that, of the potential modality decisions, only the selection of mostly-online to fully-online versus more in-person modalities (Primarily-online+), increased in likelihood as the number of cases in the county increased. This is contrary to what might be considered logical from an epidemiological perspective – whereby increased disease burden would suggest greater cause for more restrictions – and warrants further research.

The economic implications of fall instructional modality choice are best exemplified by the trends in student enrollment. With increased enrollment came more modality restriction, but only to a point – suggesting that large institutions with substantial student populations could not afford to go fully-online, either with (Fully-online+) or without (Fully-online-closed) students on campus. This finding further illustrates the increased reliance of colleges and universities on other revenue-generating activities to remain financially solvent.

In the end, there was no universally ‘right’ or ‘wrong’ decision for modality selection, as institutions had to make the best decision for themselves, considering resource availability, competing socio-political pressures, financial implications, and other factors. With the benefit of hindsight, the consequences of these decisions become clear. Perhaps the closest indicator of a right/wrong decision would be whether institutions were forced to alter operational plans mid-semester due to increased disease burden of COVID-19 or otherwise, but this is a focus for future research. Similarly, another factor to consider in future research, and related to institutional decision-making, is the background education and professional expertise (e.g. health sciences) of college and university leadership within each institution. It is still unclear as to whether the presence of public health faculty translated to their active participation in the decision-making processes, or if having a CEPH-accredited school or program merely contributed to an increased awareness and sensitivity to public health issues within and affecting the institution. If only the latter, the application of shared governance and faculty expertise may be hindered:

> “In terms of the faculty role, shared governance is just another way of saying that those with expertise in an institution’s core technology should have some important role in governing it. When that is not the case, “levels of satisfaction are likely to be low, the system may become too simple for its environment, problems are not properly attended to, and the institution may appear to lurch from crisis to crisis…”(Birnbaum, 2004, p 17)

Public health practitioners, including public health faculty, routinely make decisions on behalf of entire populations from the best available information, and with the intent to do the least harm. The Precautionary Principle remains an aspirational standard that speaks to public health professional ideals (Kriebel & Tickner, 2001). While this ecological study carries certain limitations and cannot speak to cause and effect relationships, the associations demonstrated in this study suggest that the institutional presence of public health faculty, within CEPH-accredited schools and programs, did influence colleges and universities to select less risky instructional modalities for fall 2020, and toward alternatives that consider both the individual health risks associated with COVID-19 and the consequences associated with shutting down campuses.

Institutional presence of public health in higher education is a valuable resource and can be leveraged to guide the institution toward options that honor best public health practices in a manner that allows competing priorities.

## Acknowledgements

The authors are grateful to Chris Marsicano and the research team from the College Crisis Initiative at Davidson College for the data used in this study.

## Appendix and Supplemental Materials

Equation A1. Two-level Regression

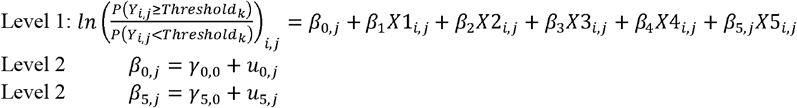

where *i* and *j* are index numbers that represent the *i^th^* college/university within each *j^th^* state *k* is an index number that represents the threshold (cut point) the divides the ordinal scale into two (in total, five thresholds are used for the 6-level modality restriction scale: (1) Levels 2-6 vs. 1; (2) Levels 3-6 vs. 1-2; (3) Levels 4-6 vs. 1-3; (4) Levels 5-6 vs. 1-4; and (5) Level 6 vs. 1-5) *β*_0,*j*_ represents the intercept for the model for each *j^th^* state, which includes both a fixed component *γ*_0,0_ (reported as the fixed intercept(s) in each model) and a component that varies by state (which is constrained to have an overall mean of 0; the variance is reported in the model results) *β*_1_ through *β*_4_ represent the fixed slopes for the CEPH accreditation, enrollment, COVID-19 rate, and AAUP variables, respectively *β*_5,*j*_ represents the slope for the Governor’s Party variable, which includes both a fixed component *γ*_5,*j*_ (reported as the fixed slope in each model) and a component that varies by state (which is constrained to have an overall mean of 0; the variance is reported in the model results)

**Table A1.**
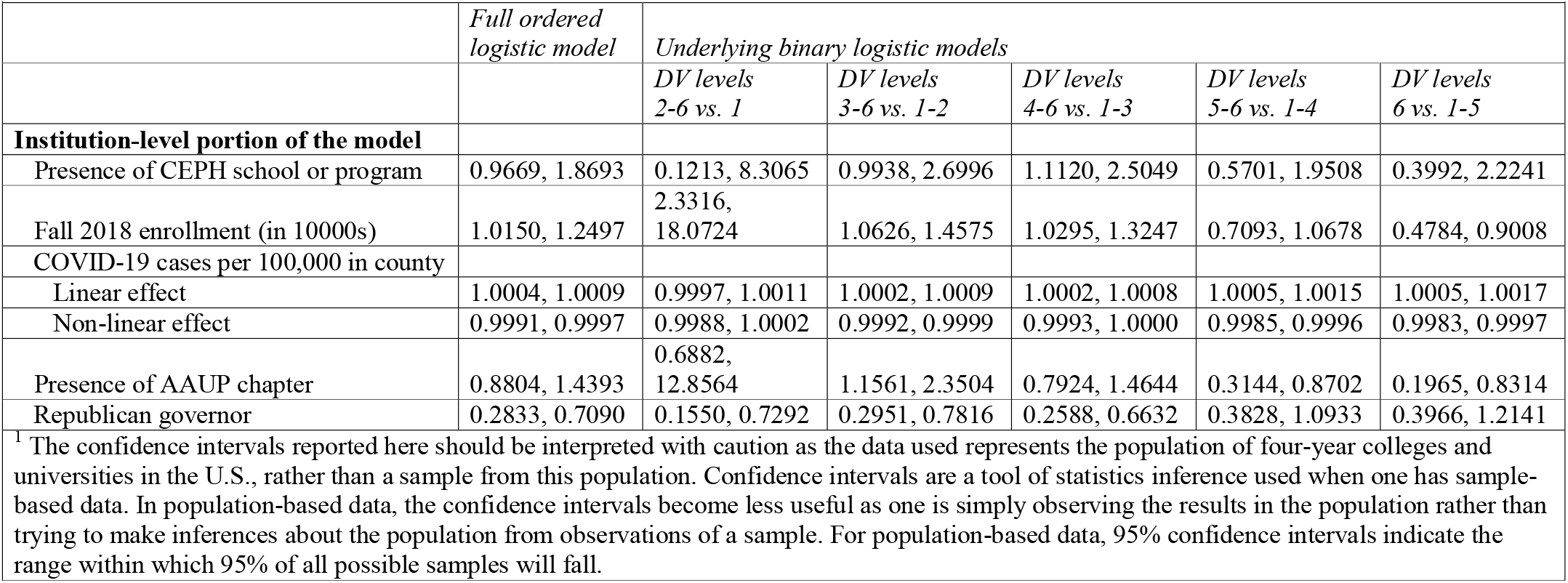
95% confidence intervals for the mixed effect logistic regression models reported in Table 2^1^

