## Appendix and Supplemental Materials 2 for "The Influence of Public Health Faculty on College and University Plans during the COVID-19 Pandemic"

| Table A1. 95% confidence intervals for the mixed effect logistic regression models reported in Table 2 ^1^ | | | | | | |
| --- | --- | --- | --- | --- | --- | --- |
|  | *Full ordered logistic model* | *Underlying binary logistic models* | | | | |
|  |  | *DV levels*  *2-6 vs. 1* | *DV levels*  *3-6 vs. 1-2* | *DV levels*  *4-6 vs. 1-3* | *DV levels*  *5-6 vs. 1-4* | *DV levels*  *6 vs. 1-5* |
| **Institution-level portion of the model** |  |  |  |  |  |  |
| Presence of CEPH school or program | 0.9669, 1.8693 | 0.1213, 8.3065 | 0.9938, 2.6996 | 1.1120, 2.5049 | 0.5701, 1.9508 | 0.3992, 2.2241 |
| Fall 2018 enrollment (in 10000s) | 1.0150, 1.2497 | 2.3316, 18.0724 | 1.0626, 1.4575 | 1.0295, 1.3247 | 0.7093, 1.0678 | 0.4784, 0.9008 |
| COVID-19 cases per 100,000 in county |  |  |  |  |  |  |
| Linear effect | 1.0004, 1.0009 | 0.9997, 1.0011 | 1.0002, 1.0009 | 1.0002, 1.0008 | 1.0005, 1.0015 | 1.0005, 1.0017 |
| Non-linear effect | 0.9991, 0.9997 | 0.9988, 1.0002 | 0.9992, 0.9999 | 0.9993, 1.0000 | 0.9985, 0.9996 | 0.9983, 0.9997 |
| Presence of AAUP chapter | 0.8804, 1.4393 | 0.6882, 12.8564 | 1.1561, 2.3504 | 0.7924, 1.4644 | 0.3144, 0.8702 | 0.1965, 0.8314 |
| Republican governor | 0.2833, 0.7090 | 0.1550, 0.7292 | 0.2951, 0.7816 | 0.2588, 0.6632 | 0.3828, 1.0933 | 0.3966, 1.2141 |
| ^1^ The confidence intervals reported here should be interpreted with caution as the data used represents the population of four-year colleges and universities in the U.S., rather than a sample from this population. Confidence intervals are a tool of statistics inference used when one has sample-based data. In population-based data, the confidence intervals become less useful as one is simply observing the results in the population rather than trying to make inferences about the population from observations of a sample. For population-based data, 95% confidence intervals indicate the range within which 95% of all possible samples will fall. | | | | | | |
